# Neural representation of nouns and verbs in congenitally blind and sighted individuals

**DOI:** 10.1101/2024.04.14.589082

**Authors:** Marta Urbaniak, Małgorzata Paczyńska, Alfonso Caramazza, Łukasz Bola

## Abstract

Language processing involves similar brain regions across languages and cultures. Intriguingly, one population escapes this universal pattern: in blind individuals, linguistic stimuli activate not only canonical language networks, but also the “visual” cortex. Theoretical implications of this finding are debated, particularly because it is unclear what properties of linguistic stimuli are represented in the blind visual cortex. To address this issue, we enrolled congenitally blind and sighted participants in an fMRI experiment, in which they listened to concrete, abstract, and pseudo nouns and verbs. We used multi-voxel pattern classification to investigate whether differences between nouns and verbs are represented in the blind visual cortex, and whether this effect is modulated by the word’s semantic category. The classification of activation patterns for nouns and verbs was above chance level in the motion-sensitive area V5/MT in the blind participants, but not in other visual areas in this group. The effect in area V5/MT was driven by successful classification of activations for concrete nouns and verbs, in the absence of significant results for abstract and pseudo nouns and verbs. These findings suggest that the blind visual cortex represents the physical properties of noun and verb referents, more salient in the concrete word category, rather than more abstract linguistic distinctions, present in all word categories. Thus, responses to language in the blind visual cortex may be explained by preserved ability of this region to compute physical and spatial representations of the world.

**Significance Statement:** In sighted individuals, language processing involves similar brain regions across languages. Intriguingly, in blind individuals, hearing words and sentences activates not only the canonical language network, but also the “visual” cortex. What is computed in the visual cortex when blind individuals process language? Here, we show that a specific visual area in the blind – the motion-sensitive area V5/MT – responds differently to spoken nouns and verbs. We further showed that this effect is present for concrete nouns and verbs, but not for abstract or pseudo nouns and verbs. This suggests that, during language processing, the blind visual cortex represents physical features of word referents, more salient in the concrete word category, rather than more abstract linguistic distinctions, present across word categories.

## Introduction

Humans acquire language relatively quickly and effortlessly. This observation has generally been taken to suggest that the human brain has strong, innate adaptations to process language. One of these adaptations might be the evolution of the brain language network - a set of brain areas that have innate capability to process complex linguistic information. Neuropsychological and neuroimaging work supports the existence of such a network by showing that the neural basis of language processing is relatively robust to changes in individual experience. For example, similar brain regions are activated by words and sentences across a variety of spoken languages and cultures (Malik-Moraleda et al., 2022), and even across spoken and sign languages (Hickok et al., 1998; MacSweeny et al., 2008). In all these cases, language processing primarily involves densely connected areas clustered around the left sylvian fissure, with the inferior frontal gyrus (Broca’s area), the superior temporal lobe (Wernicke’s area), and the anterior temporal lobe as key processing hubs (Friederici & Gierhan, 2013; Skeide & Friederici, 2016). Lesions or neurodegeneration in this network can cause a variety of language deficits, such as inability to produce or understand speech, or inability to recall object names (Hillis & Caramazza, 1991, 1995; Mesulam et al., 2014; Matchin et al., 2022).

However, the view that language can be processed only by specialized brain areas has been challenged by the observation that, in blind individuals, linguistic stimuli activate not only the canonical language network, but also the “visual” cortex (Burton et al., 2002; Bedny et al., 2011; Bedny et al., 2015; Lane et al., 2015). The magnitude of these activations increases with increasing complexity of linguistic stimuli (Bedny et al., 2011; Lane et al., 2015), and is higher for semantic than phonological tasks (Burton et al., 2003). Furthermore, a transient disruption of activity in early visual areas interferes with certain operations over linguistic stimuli in blind individuals, such as Braille reading (Cohen et al., 1997, Kupers et al., 2007) or word transformations (Amedi et al., 2004). Finally, the functional connectivity between several visual regions and the canonical language region, the left inferior frontal gyrus, is increased in blind individuals, compared to the sighted. (Bedny et al., 2011, Striem-Amit et al., 2015; Abboud et al., 2019).

The theoretical implications of these findings are still debated, particularly because it is not clear what properties of linguistic stimuli are captured by the blind visual cortex. Previous studies have convincingly shown that, in the blind, this region is activated by a variety of linguistic stimuli and tasks. However, language can be represented at various levels, from relatively concrete representation of word and sentence referents (e.g., the word “an apple” refers to an object that is small, round, etc.) to abstract representations of predominantly linguistic properties (e.g., the word “apple” is a high-frequency noun). Processing of language at these different levels can produce deceptively similar patterns of activations: A word can induce stronger activity in a given brain area than a non-word either because it names an object with certain physical properties (compare “a ball” with “an allb”) or because it is identified as a valid part of our lexicon; A sentence can induce stronger activity than a word either because it conveys more information about our physical environment (compare “a ball” with “Mary threw a ball to the dog”) or because of its syntactic structure. It is still unclear at what level linguistic stimuli can be represented in the blind visual cortex, and to what extent this representation is truly different from those computed in the visual areas of the sighted.

In this study, we addressed this issue by investigating neural representations of nouns and verbs in congenitally blind and sighted individuals. The distinction between these two word classes is rich not only in abstract grammatical consequences, but also in relatively concrete semantic implications, with many nouns naming objects and many verbs naming actions. At the brain level, both nouns and verbs seem to be represented throughout the canonical language network in sighted individuals (Ellie et al., 2019). However, verbs elicit more activity in the left frontal and posterior temporal regions, whereas nouns induce stronger responses in the left inferior temporal regions (Shapiro et al., 2005, 2006; Ellie et al., 2019). Moreover, selective impairments in production of either nouns or verbs has been demonstrated by both neuropsychological (reviewed in Shapiro and Caramazza, 2003) and neurostimulation (Shapiro et al., 2001; Cappelletti et al., 2008) studies. This shows that, in the sighted brain, the processing of these two word classes is supported by at least partly different neural representations.

We enrolled congenitally blind and sighted participants in a functional magnetic resonance imaging (fMRI) experiment, in which they made morphological transformations (singular-to-plural number transformations) to spoken concrete nouns and verbs, abstract nouns and verbs, and morphologically-marked pseudo nouns and verbs. The words were chosen based on behavioral ratings by blind and sighted participants, and task difficulty was equated across nouns and verbs (see Methods). Different words were presented in each fMRI run. We used multi-voxel pattern classification to reveal brain areas representing the noun/verb distinction across and within the three semantic categories (concrete, abstract, pseudo) in both participant groups.

We performed this study with two possibilities in mind. One possibility is that linguistic effects in the blind visual cortex are driven by, or are an extension of, typical visuospatial computations performed in this region. That would imply that the blind visual cortex represents linguistic stimuli at a relatively concrete level – one can assume, for example, that this region is sensitive to physical properties of objects and actions named by words and sentences. Based on this account, one might expect the activation patterns in the blind visual cortex to primarily reflect differences between concrete words, since a defining characteristic of these words is their having physical and spatial referents. In our study, these effects should be present in visual areas sensitive to physical properties that are differentially captured across nouns and verbs. A strong candidate for such a region is area V5/MT, which is sensitive to visual (Zeki et al., 1991) and auditory (Poirier et al., 2005; Rezk et al., 2020) motion. The auditory sensitivity of this area is elevated in blind individuals (Bedny et al., 2010; Strnad et al., 2013; Dormal et al., 2016), and can potentially be used to represent motion connotations of concrete nouns and verbs - a physical property that is differentially captured across these two word classes.

An alternative possibility is that responses to linguistic stimuli in the blind visual cortex are driven by a more fundamental form of neural plasticity. One can suppose, for example, that changes in relative strength of subcortical feedforward (Magrou et al., 2020) and cortical backward projections (Magrou et al., 2018) can lead to the development of new functional hierarchies and computational biases in this region. This can take the form of a “reverse hierarchy”, in which the low-level visual areas assume more abstract cognitive functions than the high-level visual areas, which are richly multimodal and therefore “anchored” in their typical function even in blindness (Amedi et al., 2003). Such an account opens a possibility that, at least in certain visual areas in the blind, the distinction between nouns and verbs is represented at a more abstract, conceptual or grammatical level. If this is the case, the activation patterns in these areas should capture differences between nouns and verbs from all semantic categories (concrete, abstract, and pseudo) used in the study. Based on the “reverse hierarchy” hypothesis, one might expect to find such abstract representation in the low-level visual regions in blind individuals.

## Results

### Multi-voxel pattern classification analysis

We first performed the classification of activity patterns for nouns and verbs in the visual areas when all semantic categories (concrete, abstract, pseudo) were included in one, omnibus analysis (Fig. 1). In the blind participants, this analysis produced a significant effect in area V5/MT (p = 0.008), but not in other visual areas studied (all p values > 0.14). In the sighted participants, no significant results were observed in any of the visual areas (all p values > 0.25). A direct between-group comparison confirmed that the classification of activity patterns for nouns and verbs in area V5/MT was more accurate in the blind group (t(38) = 2.94, p = 0.006).

**Figure 1.**
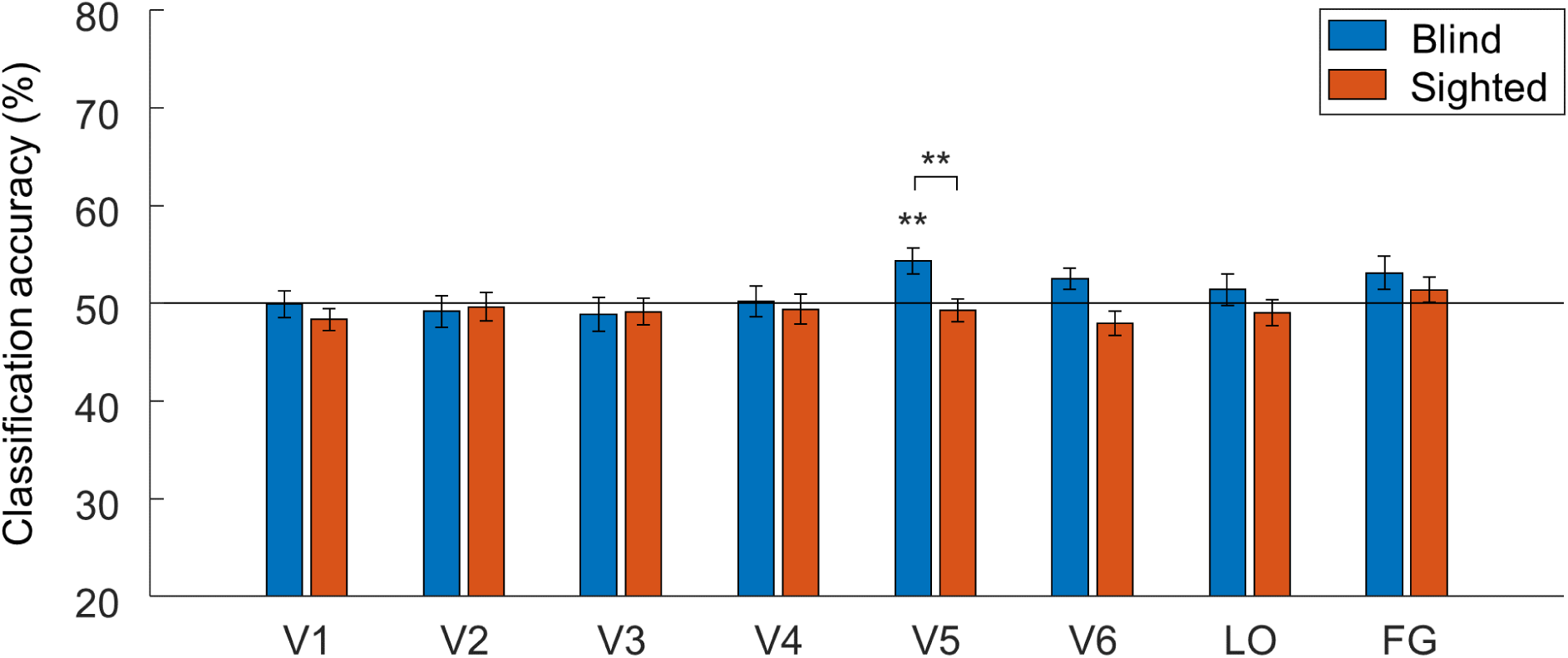
The above-chance classification of activity patterns for nouns and verbs in area V5/MT in congenitally blind individuals. Results of support vector machine classification of activity patterns for noun blocks and verb blocks in the visual areas in congenitally blind and sighted participants. LO - the lateral occipital area; FG - the fusiform gyrus. * p < 0.05, ** p < 0.01, corrected for multiple comparisons using Bonferroni correction. Error bars represent the standard error of the mean. The black line indicates the chance classification level.

We then asked whether the significant effect in area V5/MT in the blind participants could be driven by a specific word category. To address this question, we performed classifications of activity patterns for nouns and verbs in this area for each of the three word categories separately (Fig. 2). Indeed, in the blind participants, this analysis produced a significant effect for concrete nouns and verbs (p = 0.024), but not for abstract or pseudo nouns and verbs (both p values > 0.25). The classification accuracy for concrete nouns and verbs was significantly higher than the average classification accuracy calculated across the abstract and pseudo word categories (t(19) = 2.48, p = 0.023). In the sighted participants, the same analysis did not produce significant results in any word category (all p values > 0.25). However, a 2 (group) x 3 (word category) ANOVA indicated a significant main effect of word category (F(2,76) = 3.42, p = 0.038, partial eta-squared = 0.08), with no main effect of group and no interaction between these two factors (both p values > 0.25). Thus, while the ANOVA confirmed the differences in the classification of activity patterns for nouns and verbs across the three word categories in area V5/MT, this analysis did not provide strong evidence that this effect is specific to the blind group.

**Figure 2.**
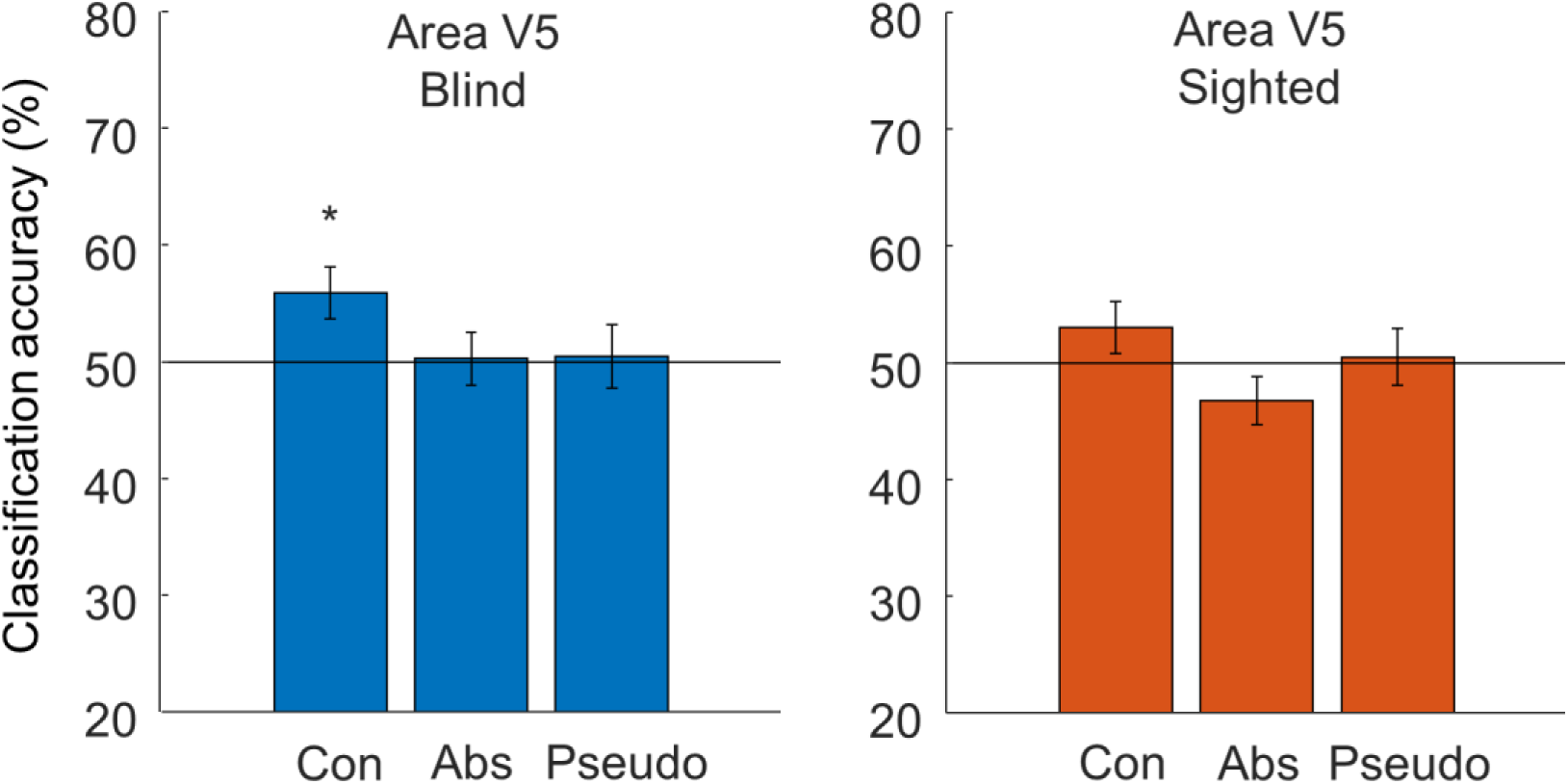
The effect in area V5/MT in blind individuals is driven by successful classification of activity patterns for concrete nouns and verbs. Results of support vector machine classification of activity patterns for noun blocks and verb blocks, performed separately for concrete, abstract, and pseudo word categories, in area V5/MT in congenitally blind and sighted participants. * p < 0.05, corrected for multiple comparisons using Bonferroni correction. Error bars represent the standard error of the mean. The black lines indicate the chance classification level.

As a control analysis, we investigated the accuracy of classification of activity patterns for nouns and verbs for each word category in all visual areas considered in the study (Fig.S1). This analysis was meant to search for meaningful effects that were specific to only one word category and could be potentially missed in the initial, omnibus analysis. However, apart from the already reported effect in area V5/MT in the blind participants, we only observed above-chance classification of activations for pseudo nouns and verbs in several visual areas in the blind participants. This result could perhaps be driven by some form of surprise response to these atypical stimuli. Critically, this effect was not accompanied by the successful classification of activations for abstract nouns and verbs in any of the visual areas (all p values > 0.25). Furthermore, besides the already reported effect in area V5/MT, the classification of activations for concrete nouns and verbs did not produce significant results in any other visual area (all p values > 0.25). This confirms the topographic specificity of the effect reported for concrete nouns and verbs in area V5/MT.

Next, we investigated what brain regions, beyond the visual cortex, capture the differences between nouns and verbs in the blind and the sighted participants. To this aim, we performed an omnibus classification of activity patterns for nouns and verbs, with words from all three categories included, using a whole-brain searchlight approach (Fig. 3). In both participant groups, we observed significant effects primarily along the superior temporal sulci (Fig. 3A-B). In the blind participants, we additionally detected significant effects in the left inferior frontal gyrus (Brodmann areas 44 and 45), the left fusiform gyrus, and the left insula. A direct between-group comparison confirmed that the results in the opercular part of the left inferior frontal gyrus (Brodmann area 44) and in the insula were stronger in the blind group (Fig. 3C). There were no effects that were stronger in the sighted group.

**Figure 3.**
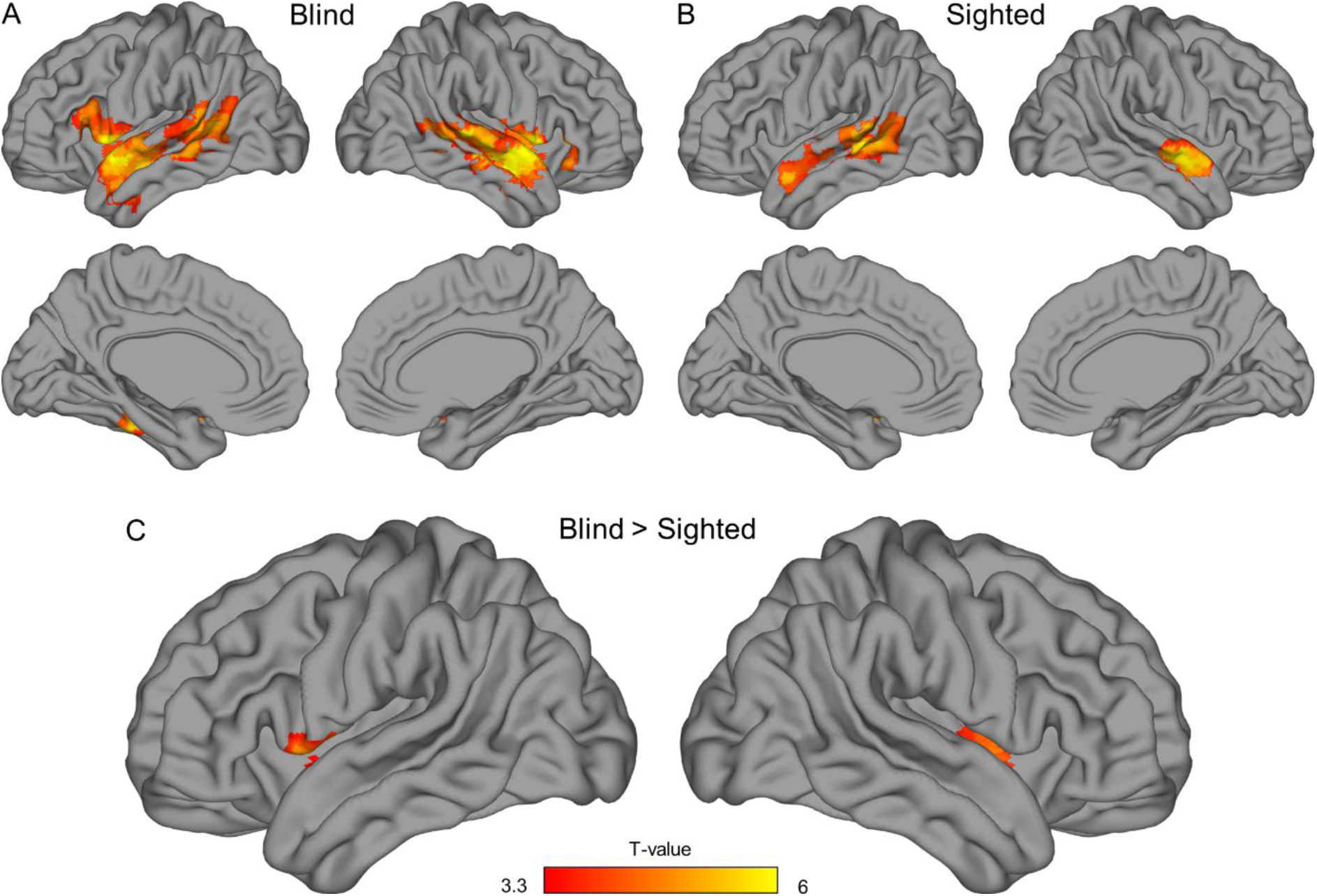
A searchlight classification of activations for nouns and verbs. Results of support vector machine classification of activity patterns for noun blocks and verb blocks, performed using a searchlight procedure (A) in congenitally blind participants, (B) in sighted participants, and (C) when blind participants were compared to the sighted. Statistical threshold was set to p < 0.001, corrected for multiple comparisons using a family-wise error cluster correction.

We further asked whether the superior temporal cortex – the canonical language region in which the omnibus analysis produced the strongest results in both groups – captures the differences between nouns and verbs for all three word categories included in the study. To answer this question, we performed an independent region of interest analysis in this area (Fig. 4). Indeed, we found above-chance classification of activations for nouns and verbs from all three word categories in both the blind and the sighted participants (classification of concrete nouns and verbs in the blind group: p = 0.057; all other p values < 0.05), with no significant differences across the word categories in either of the participant groups (all p values > 0.25). A 2 (group) x 3 (word category) ANOVA did not produce any significant main effects or interactions (all p values > 0.25). This shows that the specific effect observed for concrete nouns and verbs in area V5/MT could not be explained by better classification of concrete words throughout the brain.

**Figure 4.**
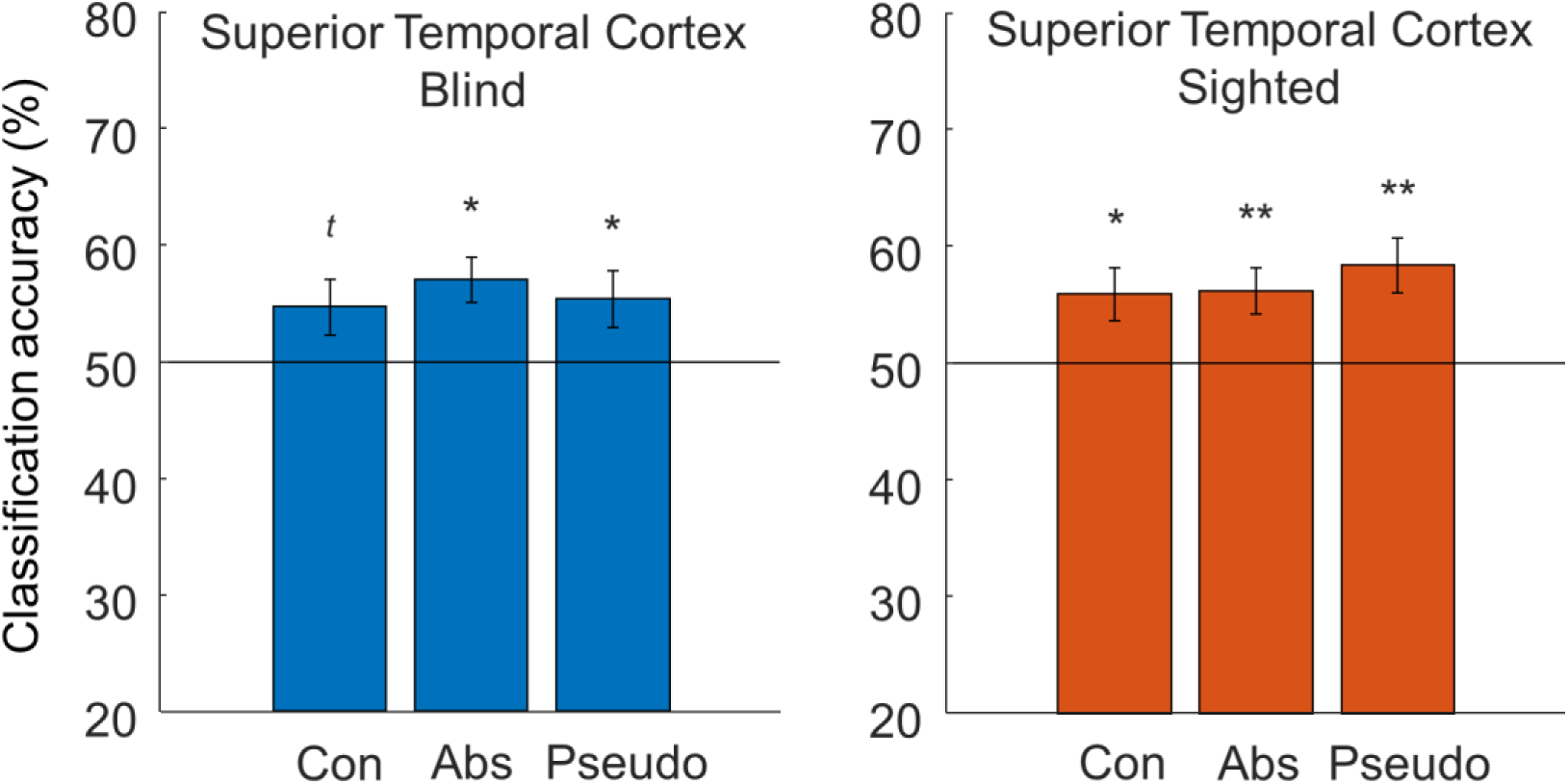
The effects in the superior temporal cortex are driven by successful classification of activity patterns for nouns and verbs from all semantic categories. Results of support vector machine classification of activity patterns for noun blocks and verb blocks, performed separately for concrete, abstract, and pseudo word categories, in the superior temporal cortex in congenitally blind and sighted participants. * p < 0.05, ** p < 0.01, *^t^* p = 0.057, corrected for multiple comparisons using Bonferroni correction. Error bars represent the standard error of the mean. The black lines indicate the chance classification level.

Additionally, we directly tested whether the result patterns in the superior temporal cortex and area V5/MT were different. Indeed, a 2 (brain area) x 2 (group) x 3 (word category) ANOVA produced a significant main effect of area (F(1,38) = 19.077, p < 0.001, partial eta-squared = 0.33) and a trend-level interaction between the area and the word category factors (F(2,76) = 2.67, p = 0.076, partial eta-squared = 0.07), with no other main effects or interactions being significant (all other p values > 0.15). The post hoc tests indicated that, compared to area V5/MT, the classification accuracy in the superior temporal cortex was significantly higher for the abstract (p < 0.001) and pseudo (p = 0.008) nouns and verbs, but not for concrete nouns and verbs (p > 0.25).

Finally, the hypothesis that the blind visual cortex is sensitive to words because it represents the physical features of word referents implies that this region captures differences not only between concrete nouns and verbs, but also between concrete and abstract words. While our design was not optimized for this contrast, we tested this prediction and found a robust, above-chance classification of activity patterns for concrete and abstract words in areas V4 (p = 0.018) and V5 (p = 0.027) in the blind participants (Fig. S2). In contrast, the classification of activations for concrete and abstract words was not successful in the superior temporal cortex, in any of the participant groups (both p > 0.25) (Fig. S2), which further suggests that the effects observed in this canonical language region might be driven by different neural representation than the effects found in the visual cortex in blind individuals.

### Univariate analysis

We also performed univariate analyses to verify whether the visual cortex in blind participants responded to spoken words more strongly than the visual cortex in sighted participants. The whole-brain analysis (Fig. 5) showed that, in both groups, listening to spoken words and pseudowords activated the canonical language regions (Fig. 5A-B). In the blind participants, significant responses to these stimuli were also observed in the high-level visual areas. The direct between-group comparison showed that both early and high-level visual areas responded to spoken words more strongly in the blind group than in the sighted group (Fig. 5C). An additional region of interest analysis (Fig. S3) showed that this effect was driven by both activations in the visual cortex in the blind participants and deactivations in the visual cortex in the sighted participants.

**Figure 5.**
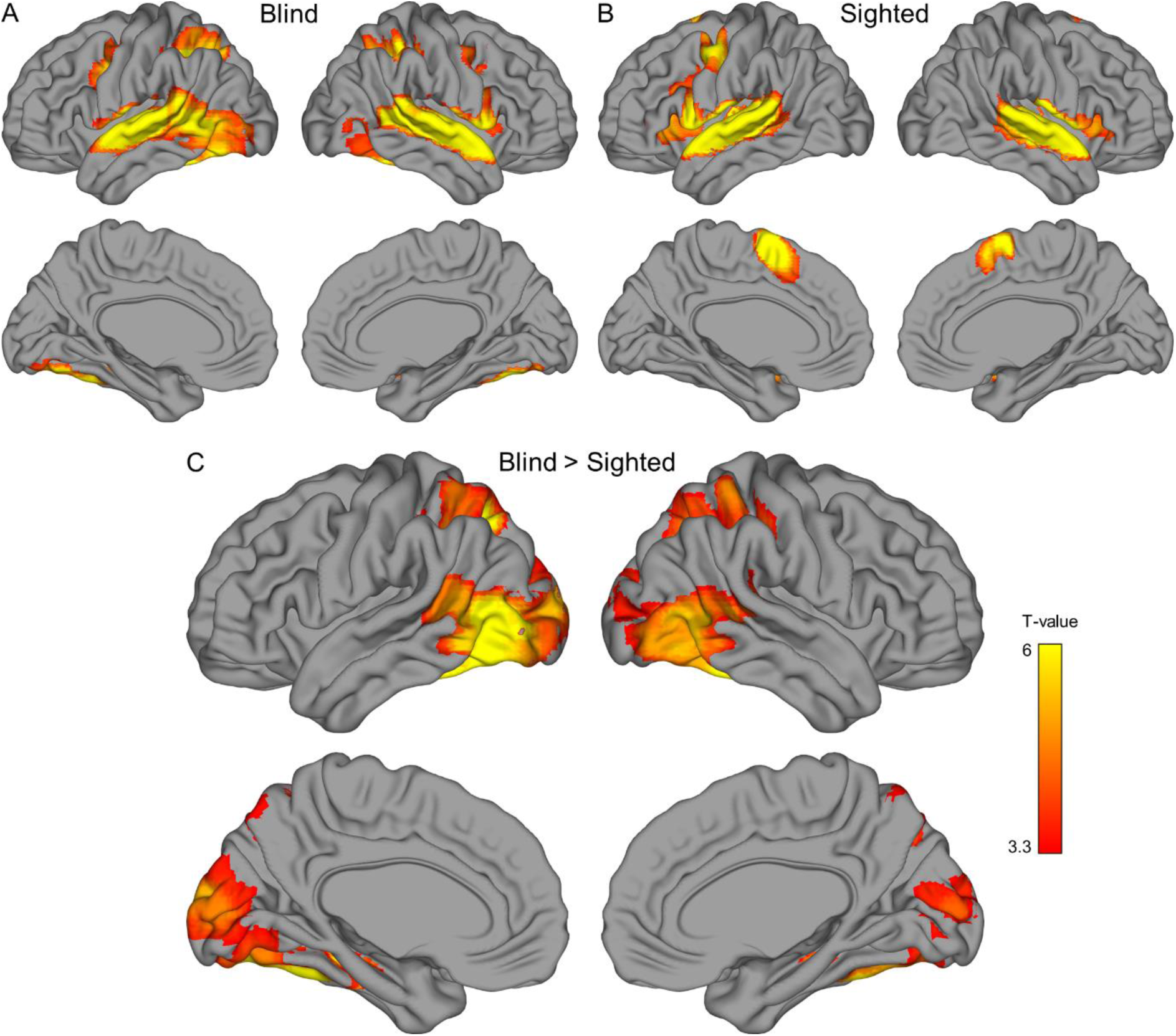
Brain activations for spoken words in congenitally blind and sighted individuals. Average responses to all spoken words and pseudowords, compared to rest periods, (A) in congenitally blind participants, (B) in sighted participants, and (C) when the blind participants were compared to the sighted. Statistical threshold was set to p < 0.001, corrected for multiple comparisons using a family-wise error cluster correction.

Finally, we investigated univariate activations elicited by specific word classes in area V5/MT (Fig. 6), in which multi-voxel activation patterns were different for concrete nouns and verbs. In the blind participants, we observed significant activation of this area for all word classes (all p values < 0.05). In the sighted participants, in contrast, all word classes induced significant deactivation of this region, relative to rest periods (all p values < 0.05). A 2 (group) x 2 (grammatical class) x 3 (word category) ANOVA indicated a significant main effect of group (F(1,38) = 37.36, p < 0.001, partial eta-squared = 0.5), a significant main effect of word category (F(2,76) = 4.16, p = 0.019, partial eta-squared = 0.1), and interaction between these two factors (F(2,76) = 18.21, p < 0.001, partial eta-squared = 0.32). Interestingly, neither main effect of grammatical class nor interactions including this factor were significant (all p values > 0.2). Pairwise comparisons showed that, in the blind participants, pseudowords induced stronger activation in area V5/MT than concrete words or abstract words (both p values < 0.001). In contrast, no differences across word categories were observed in the sighted participants (all p values > 0.05). There were no differences between nouns and verbs in any word category or participant group (all p values > 0.05). These results suggest that global, univariate responses might capture different neural processes in area V5/MT of blind individuals than the multi-voxel activation patterns.

**Figure 6.**
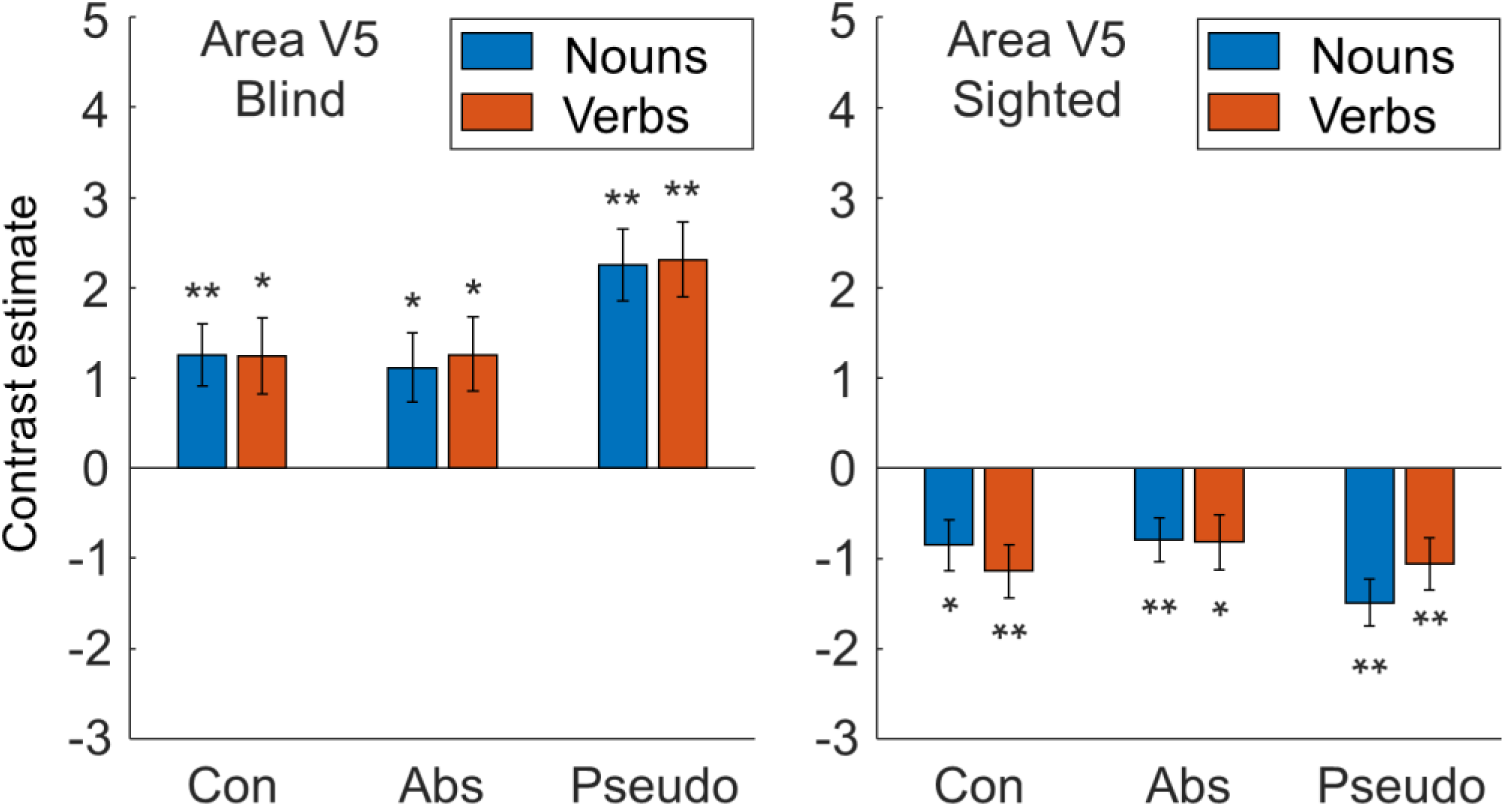
Brain responses to specific word categories in area V5/MT in congenitally blind and sighted individuals. The responses are presented relative to activations during rest periods. * p < 0.05, ** p < 0.01, corrected for multiple comparisons using Bonferroni correction. Error bars represent the standard error of the mean.

## Discussion

In this study, we found that classification of activity patterns for nouns and verbs was significantly above chance level in area V5/MT in congenitally blind participants, but not in other visual areas in this group. We further showed that the effect in area V5/MT in the blind was driven by successful classification of activations for concrete nouns and verbs, in the absence of significant results for abstract and pseudo nouns and verbs. Beyond the visual cortex, we found successful classification of activity patterns for nouns and verbs primarily in the superior temporal cortex, in both blind and sighted participants, with additional effects in several other canonical language regions in the blind. In both groups, the activity patterns in the superior temporal cortex captured differences between nouns and verbs from all three word categories (concrete, abstract, pseudo) used in the study.

In this study, we asked what properties of linguistic stimuli are represented in the visual cortex of congenitally blind individuals, and can drive the activations of this region for language (Burton et al., 2002; Burton et al., 2003; Bedny et al., 2011; Lane et al., 2015). We focused on a fundamental linguistic distinction – between nouns and verbs – and investigated what aspects of this distinction are represented in the blind visual cortex. We found above-chance classification of activity patterns for nouns and verbs in the visual area V5/MT in the blind participants, but not in other visual areas in this group. We further showed that the effect in area V5/MT in the blind was driven by successful classification of activations for concrete nouns and verbs, in the absence of significant results for abstract and pseudo nouns and verbs. Area V5/MT is sensitive to visual (Zeki et al., 1991) and auditory (Poirier et al., 2005, Rezk et al., 2020) motion, with the auditory sensitivity being preserved and elevated in blind individuals (Bedny et al., 2010; Strnad et al., 2013; Dormal et al., 2016). Here, we suggest that this area’s functional specialization for motion processing can be used to represent motion connotations conveyed by concrete nouns and verbs. This physical property is strongly conveyed by concrete words, but not necessarily by other word categories used in the study. Crucially, it is also differently captured by concrete nouns (generally naming stationary objects) and concrete verbs (generally naming dynamic actions), which can lead to different activity patterns induced in area V5/MT by these two word classes. This effect is more robust in the blind individuals presumably because of the elevated sensitivity of this area to auditory signals in this population.

Our findings are in line with demonstrations that, in both blind and sighted individuals, high-level ventral visual regions respond differently to words referring to objects of different shape and size (Mahon et al., 2009; He et al., 2013; Peelen et al., 2013, 2014). Similar effects were shown in these regions during the presentation of object pictures (e.g., Konkle and Caramazza, 2013; Bracci and Op de Beeck, 2015; Long et al., 2018). Here, we show that also the dorsal visual regions can use information conveyed by spoken words to perform their relatively typical computations, such as representation of motion and motion connotations. Crucially, we used abstract and pseudo words to test for more abstract, linguistic representation in the blind visual cortex, as such representation could be potentially computed on top of simpler representations of physical properties. However, we did not find any clear sign of such abstract representation in any visual area tested.

Overall, our results indicate that the blind visual cortex represents the physical properties of word referents, more salient in the concrete word category, rather than more abstract linguistic features, present across the word categories. The topography of effects observed in our study suggests that these physical connotations, conveyed by words, are mapped onto the typical functional organization of the visual cortex, present also in the sighted brain.

Our findings contribute to a better understanding of principles of plasticity in the human brain. One way to think of language-driven activation in the blind visual cortex is that, in the absence of visual signals, this region develops new functional properties, which are radically different from those computed in the sighted visual cortex (Amedi et al., 2003; Bedny et al., 2017). Our findings provide a different perspective and suggest that at least some effects observed for language in the blind visual cortex might be explained by typical computational biases and preserved ability of this region to compute physical and spatial representations of the world. In the visual cortex of sighted individuals, the visuospatial representation emerges in interaction between feedforward pathways and modulatory back projections from higher-order brain regions. In blind individuals, many of these back projections are known to be preserved (Magrou et al., 2018) and functional (Vetter et al., 2020; Bola et al., 2023), and perhaps can communicate physical properties of objects and actions, conveyed by spoken words, to the visual areas. In this view, responses to linguistic stimuli in the blind visual cortex can be driven by strengthening and uncovering of functional interactions between higher-order and visual cortices that are present also in the sighted brain (Pascual-Leone and Hamilton, 2001).

The univariate analyses reported here showed activations for spoken words in the visual cortex in blind participants, a result that is consistent with previous studies. Interestingly, the analysis in area V5/MT in this group suggests that the global magnitude of activation in this region, as measured by univariate methods, captures different processes than the multi-voxel activation patterns. The global activation of area V5/MT in the blind was the same for all word classes, but higher for pseudowords, which suggests that this measure captured increased processing demands related to surprising or atypical stimuli. Attention or cognitive load modulates the activation magnitude in the visual cortex even in sighted individuals (Gandhi et al., 1999; Watanabe et al, 2011). The weakening of inhibitory mechanisms in the blind visual cortex (Benevento et al., 1995; Morales et al., 2002; Keck et al., 2011) might strengthen these effects, leading to the generic response that can cover other processes, specific for a given word category. These more specific processes can be nevertheless detected with more sensitive multivariate analysis.

Conversely, in the visual cortex of sighted individuals, spoken words induced robust deactivation, relative to the rest periods. This effect might be driven by a known mechanism of inhibition of the visual system activity during auditory perception (Laurienti et al., 2002; Iurilli et al., 2012; Anurova et al., 2019). This process might be less efficient in blind individuals due to generally weaker inhibition mechanisms in their visual areas, described above (see also Anurova et al., 2019). One study demonstrated that certain regions within the early visual cortex of sighted individuals are activated by spoken words (Seydell-Greenwald et al., 2023). Here, we did not find such an effect. One explanation could be the differences in tasks used in the two studies. Seydell-Greenwald and colleagues used passive listening of spoken words. In this study, we used a relatively demanding morphological transformation task. This higher cognitive demand could potentially strengthen deactivation of the visual cortex in sighted individuals.

Beyond the visual cortex, we observed the representation of differences between nouns and verbs only in the canonical language regions, in both the blind and the sighted participants. While, in both groups, this representation was strongest in the middle and superior temporal cortices, there were several, additional effects in other canonical language regions in the blind participants. One possible explanation is that blind individuals are more attentive to speech (Hugdahl et al., 2004), which can lead to more robust signal in certain language areas. Despite these between-group differences, our results suggest that the overall topography of the canonical language network is robust to changes in visual experience.

In conclusion, our findings suggest that the blind visual cortex represents the physical properties of word referents rather than more abstract, conceptual or grammatical distinctions. This shows that at least some effects observed for language in the blind visual cortex might be explained by preserved ability of this region to compute physical and spatial representations of the world. The topography of effects observed in our study suggests that, in blind individuals, physical connotations conveyed by spoken words are mapped onto the typical functional organization of the visual cortex, present also in the sighted brain.

## Methods

### Participants

Twenty congenitally blind subjects (9 males, 11 females, mean age = 35.65 y, SD = 7.81 y, average length of education = 14.8 y, SD = 2.35 y) and 20 sighted subjects (6 males, 14 females, mean age = 35 y, SD = 8.58 y, average length of education = 15.4 y, SD = 2.04 y) participated in the study. All except two participants were right-handed, and the remaining two participants (one blind, one sighted) were left-handed. The blind and the sighted groups were matched for age, gender, handedness, and years of education (Mann-Whitney and Chi-square tests, all p values > 0.25). In the blind group, blindness had a variety of causes, including retinopathy of prematurity, glaucoma, Leber’s congenital amaurosis, optic nerve hypoplasia, or unknown causes. Most blind participants reported to have some light perception, but no object or contour vision. One blind participant reported to have some form of contour vision, which, however, was not precise enough to be functional. All subjects in both groups were native Polish speakers, had normal hearing, and had no history of neurological disorders. All subjects had no contraindications to the MRI, gave written informed consent and were paid for participation. The study was approved by the ethics committee of Institute of Psychology, Polish Academy of Sciences.

### Stimuli

In total, 144 stimuli were used: 24 concrete nouns, 24 concrete verbs, 24 abstract nouns, 24 abstract verbs, 24 pseudo nouns, and 24 pseudo verbs. All stimulus categories were matched on average number of syllables, and all words categories were additionally matched on average frequency of occurrence in Polish language (quantified as Zipf score; van Heuven et al., 2014), as indicated by Subtlex-pl database (Mandera et al., 2014) (Table S1). The length and the frequency matching were performed not only for the stimulus forms heard by the participants, but also for the target forms that the participants were expected to produce in the fMRI (Kruskal-Wallis tests, all p vaules > 0.25). All chosen words had to fulfill semantic criteria – broadly, all concrete words referred to objects or actions that are well specified in space (e.g., “a cup” or “to kick”), whereas all abstract words referred to concepts or conceptual actions without a clear spatial framework (e.g., “fairness” or “to think”; see the Appendix 1 for English translations of all words used in the study). All pseudowords were phonologically and grammatically valid but had no meaning in Polish. The pseudowords were created by mixing syllables taken from the actual words. The stimuli were audio recorded using speech synthesizer software. The recordings were judged as sounding natural and readily understandable by Polish native speakers during pilot studies.

### Behavioral experiments

The final stimuli were chosen from a larger initial dataset (240 items, 40 per category) based on the results of two pilot behavioral studies. In the first study, 16 sighted participants (8 males, 8 females, mean age = 27.5 y, SD = 7.69 y, average length of education = 14.86 y, SD = 2.94 y) were asked to transform the heard words and pseudowords, from singular to plural form, based on the verbal transformation cues, the same as were used in the fMRI experiment (see the “fMRI experiment” section below). Each item was repeated two times (480 trials in total) and the subjects were asked to produce an overt response, which was recorded using a microphone. Next, response times for each item (from the onset of word presentation to the onset of response) were calculated using Chronoset (Roux et al., 2017). The response times across stimulus categories were matched as well as possible. In the final stimulus list, response times were matched across all word categories (a Kruskal-Wallis test, p > 0.25) and across pseudo nouns and pseudo verbs (a Mann-Whitney test, p > 0.25) (see Tables S2 for average response times for all stimulus categories in the final stimulus list). As could be expected, the response times for pseudo nouns and pseudo verbs were higher than for any word category (Mann-Whitney tests, all p values < 0.05).

In the second study, 15 congenitally blind (5 males, 10 females, mean age = 36.27 y, SD = 7.89 y) and 46 sighted participants (23 males, 23 females, mean age = 25.46 y, SD = 7.69 y) were asked to rate each word from the initial list on three scales (from 1 to 7): concreteness, imaginability, and movement connotations. Then, the items with unexpected ratings (e.g., concrete words with relatively low concreteness scores) were excluded. In the final stimulus set, all word categories were rated as expected by both groups – that is, concreteness and imaginability scores were higher for the concrete words than for the abstract words (Mann-Whitney tests, all p values < 0.001), and movement connotation scores were higher for verbs, particularly the concrete ones, than for nouns (a Mann-Whitney test, p < 0.001) (see Tables S3-4 for average rating scores for all stimulus categories in the final stimulus list). Notably, there was a very high correlation between ratings provided by congenitally blind and sighted subjects, both in the initial and in the final stimulus list (Pearson’s r values > 0.9, p values < 0.001, for all three scales).

### fMRI experiment

In the fMRI experiment, the participants heard words and pseudowords in singular forms and were asked to mentally (i.e., without an overt response) transform them into plural forms, based on the verbal transformation cue presented beforehand (“many” and “few” for nouns, “we” and “they” for verbs). The subjects were explicitly instructed to treat pseudowords as the “real words” and produce transformations that sounded correct. Each word/pseudoword presentation lasted approximately 0.5 s and was followed by 2 s of silence during which subjects were asked to create a correct word form in their minds. The time assigned for each trial was set at duration well above the average response times obtained in the pilot behavioral experiment (see Table S2), to ensure that the participants were able to complete the task successfully.

The stimuli were presented in blocks of 6 items belonging to the same category (e.g., 6 abstract nouns), resulting in blocks lasting 15 s. The blocks were further grouped into “super blocks” according to the word grammatical class, such that 3 noun blocks (one with abstract nouns, one with concrete nouns, and one with pseudo nouns) were always grouped together and followed by 3 verb blocks, and vice versa. This second-level order was introduced to save time assigned for the presentation of transformation cues (i.e., the cue was the same for all blocks from the same grammatical class – thus, the introduction of the super blocks allowed us to present the cue only once per three blocks). Each super block started with a transformation cue, which was followed by 12 s of silence. Then, the three noun or three verb blocks were presented, each followed by 12 s of silence. Subsequently, the super block for the other grammatical class started. There were 4 noun super blocks and 4 verb super blocks in each fMRI run, resulting in presentation of 4 blocks for each stimulus category in each run. Subjects completed 4 runs, each lasting approximately 12 mins and 45 s.

Each run involved presentation of different words and pseudowords (6 items per category per run). Thus, all between-run decoding analyses that we performed could not rely on word repetitions, and instead had to rely on a more abstract representation of linguistic properties of specific word/pseudoword classes. In each run, the same words were repeated in each block belonging to a specific experimental condition, but the presentation order was randomized. Furthermore, the block order, within each super block, was randomized with a constraint that the same order was applied to noun super blocks and verb super blocks.

In Polish, word grammatical classes have systematically different suffixes, which might result in certain analyses of differences between nouns and verbs being confounded by phonology. Furthermore, one can assume that the phonology of the verbal transformation cues, which subjects are likely to keep in mind during transformations, can confound these results. To be able to control for these issues, we systematically varied the phonology of nouns, verbs, and transformation cues across the odd and even fMRI runs. In one type of runs we used “many” and “we” transformation cues for nouns and verbs, respectively, whereas in the other type of runs the transformation cues were “few” and “they”. Different transformation cues resulted in different inflections for verbs, but not necessarily for nouns. Thus, in the case of nouns, one type of runs additionally included only masculine nouns, whereas the other type of runs included only feminine and neutral nouns. These manipulations resulted in a design in which any decoding analysis in an odd-even cross-validation scheme could not be driven by phonology. The run order was randomized across subjects, keeping the odd-even scheme in place.

The stimuli presentation was controlled by a program written in PsychoPy 3.0.12b (Peirce, 2007). The sounds were presented through MRI-compatible headphones. Before starting the experiment, each participant completed a short training session. Furthermore, the volume of sound presentation was individually adjusted to a loud, but comfortable level. The sighted participants were blindfolded for the duration of the fMRI experiment to create as similar environment of data acquisition for the sighted and the blind group as possible.

### Imaging Parameters

Data were acquired on a 3-T Siemens Trio Tim MRI scanner using a 32-channel head coil at the Laboratory of Brain Imaging in Nencki Institute of Experimental Biology in Warsaw. Functional data were acquired using a multiband sequence with the following parameters: 60 slices, phase encoding direction from posterior to anterior; voxel size: 2,5 mm^3^; TR = 1.41 s; TE: 30.4 ms; multiband factor: 3. Before the start of the first functional run, T1-weighted anatomic scans were acquired using MPRAGE sequence with the following parameters: 208 slices, phase encoding direction from anterior to posterior; voxel size: 0,8 mm^3^; TR = 2.5 s; TE: 21.7 ms.

### MRI data analysis

#### Data preprocessing

The MRI data were converted from the DICOM format to the NIFTI format using the dcm2niix (Li et al., 2016). Then, the preprocessing was performed using SPM 12 (Wellcome Imaging Department, University College, London, UK, http://fil.ion.ucl.ac.uk/spm) and CONN 21b toolbox (Whitfield-Gabrieli & Nieto-Castanon, 2012) running on MATLAB R2022a (MathWorks Inc. Natick, MA, USA). Data from each subject were preprocessed using the following routines: (1) functional realignment of all functional images; (2) direct segmentation and normalization, where functional and anatomical data were segmented into gray matter, white matter, and CSF tissue classes and normalized into standard Montreal Neurological Institute (MNI) space using unified segmentation and normalization procedure (Ashburner & Friston, 2005), and (3) spatial smoothing with Gaussian kernel at 8-mm FWHM for the univariate analysis; no spatial smoothing for the multi-voxel pattern classification analysis.

Two first-level statistical models were created for each subject. For the multi-voxel pattern classification analysis, the data were modeled at the level of single blocks (24 predictors per run, one for each block). Additionally, transformation cues were modeled as conditions of no interest (8 predictors per run, one for each occurrence of the cue). For the univariate analysis, the data were modeled at the level of word categories (6 predictors per run, one for each word category, 1 predictor of no interest per run for cues). Signal time course was modeled using a general linear model (Friston et al., 1995) by convolving a canonical hemodynamic response function with the time series of predictors. Six movement parameter regressors obtained during the preprocessing were added to the models. An inclusive high-pass filter was used (378 s, approximately 2 cycles per run) to remove drifts from the signal while ensuring that effects specific to each word category were not filtered out from the data. Finally, individual beta maps, contrast maps, and t-maps were computed for each experimental block/condition, relative to rest periods.

#### Multi-voxel pattern classification

All multi-voxel pattern classification analyses were performed in CosmoMVPA (v.1.1.0; Oosterhof et al., 2016), running on Matlab R2022a (MathWorks). The analyses were performed on contrast maps for specific experimental blocks compared to rest periods (24 maps per run, 96 maps per participant in total). A linear support vector machine classification algorithm was used, as implemented in the LIBSVM toolbox (v. 3.23; Chang and Lin, 2001). A standard LIBSVM data normalization procedure (i.e., Z-scoring beta estimates for each voxel in the training set and applying output values to the test set) was applied to the data before classification. The region of interest (ROI) analyses were performed using maps from the JuBrain Anatomy Toolbox (Eickhoff et al., 2005). All ROIs were defined bilaterally.

We first performed the omnibus ROI classification of activations for noun blocks and verb blocks in the visual areas, without dividing the stimuli into concrete, abstract, and pseudo words. The analysis included all occipital and occipitotemporal regions delineated in the JuBrain Anatomy Toolbox. To reduce a number of tests, subregions were combined (e.g., areas V3d and V3v were combined in area V3). The classification was performed in an odd-even cross validation scheme, that is, the classifier was trained on the odd runs and tested on the even runs, and vice versa, resulting in two cross validation folds. This scheme was used to ensure that the representation of phonological information did not affect the results (see also the “fMRI Experiment” section). The results were corrected for multiple comparisons across all ROIs included in the analysis, for each participant group separately (see below for the details on statistical thresholds).

We further investigated the pattern of results in area V5/MT, in which the omnibus analysis showed significant effects. To this aim, we performed the classification of activations for noun blocks and verb blocks in this area, separately for each word category (concrete, abstract, pseudo). This more detailed analysis was performed in a leave-one-run-out cross validation scheme, that is, the classifier was iteratively trained on all runs except one and tested on the remaining run, resulting in four cross validation folds. This scheme let us test our hypothesis with maximal statistical power; at the same time, we expected positive results only for concrete word category, which means that the remaining word categories could serve as a control for phonological representation. The results were corrected across the three word categories, for each participant group separately.

We then performed the searchlight analysis to investigate which brain regions, beyond the visual cortex, capture differences between nouns and verbs in both participant groups. To this aim, we again used the omnibus classification approach, in which the experimental blocks were classified into noun blocks and verb blocks, without dividing the stimuli into concrete, abstract, and pseudo words. The classification was performed in volume space, in searchlight spheres with 5-voxel radius, using the odd-even cross validation scheme, similarly as the initial, omnibus ROI analysis.

We further studied the pattern of results in the superior temporal cortex, which showed the strongest effects in the searchlight analysis. We used a map of area TE3 from the JuBrain Anatomy Toolbox as a proxy of the superior temporal cortex, to define the ROI independently of the searchlight analysis. We the classified activity patterns for noun blocks and verb blocks in this region, separately in each word category. The analysis was performed and corrected in the same way as the category-specific analysis in the visual area V5/MT.

Finally, as a supplementary analysis, we also performed the classification of activity patterns for concrete word blocks and abstract word blocks. The classification was performed in a leave-one-run-out cross validation scheme, as the phonology of words was not systematically different across the concrete and abstract word categories. The classification was performed in both the visual ROIs and the superior temporal cortex ROI. The results were corrected across all ROIs included in the analysis, for each participant group separately.

In all ROI classification analyses, the statistical significance of obtained classification accuracies was tested against chance levels that were empirically derived in the permutation procedure. Specifically, each classification analysis was re-run 1000 times for each participant with the labels of classified conditions randomly assigned to experimental blocks in each iteration. Null distributions created in this procedure were averaged across participants and compared with the actual average classification accuracies. The p values that were obtained in this way were corrected for multiple comparisons, as was described above, using the Bonferroni correction. A review of null distributions confirmed that, for each ROI and analysis, the empirically-derived chance levels were indistinguishable from a priori chance levels (50%). Thus, for simplicity, the a priori chance level is presented in the figures.

Testing for differences in classification accuracy between the participant groups was performed with two-sample t-test. These tests were run only for ROIs in which successful classification was observed for at least one participant group. Testing for differences in classification accuracy between two ROIs or conditions was performed with a paired t-test. Testing for significant interactions between the results for ROIs, conditions, and participant groups was performed with mixed ANOVAs. SPSS 25 (IBM Corp, Armonk, NY) was used to perform these tests. Bonferroni correction was used to correct the results for multiple comparisons.

In the searchlight analysis, the individual classification results were entered into SPM group models. One-sample t-tests were used to compare the results of each searchlight analysis with chance level, separately in the blind and the sighted group. Two-sample t-tests were used to compare the results between groups. A statistical threshold for all group analyses was set at p < 0.001 voxel-wise, corrected for multiple comparisons using family-wise error cluster (FWEc) correction approach.

#### Univariate analysis

We first performed the whole-brain univariate analysis, in which we compared the activation induced by all words and pseudowords to rest periods, in both participant groups. The activations for all experimental conditions, relative to rest periods, were averaged at the single-subject level. Then, the average activation maps were entered into SPM one-sample t-tests, performed separately for each group, and into the SPM two-sample t-test, which tested for the between-group differences. As in the searchlight classification analysis, the statistical threshold for these analyses was set at p < 0.001 voxel-wise, corrected for multiple comparisons using FWEc correction approach.

The whole-brain analysis was followed by the ROI analysis in the visual cortex. The analysis of activations for all experimental conditions, relative to rest periods, was performed in the same visual ROIs that were used in the multi-voxel pattern classification analysis. Then, one-sample t-tests were used to compare the activations for all conditions with activations during rest periods, separately in the blind and the sighted group. Bonferroni correction was used to correct the results for multiple comparisons across all ROIs used in the analysis, for each participant group separately. Two-sample t-tests were then used to compare the results between groups, within each ROI that showed significant results in at least one participant group.

Finally, we performed the ROI analysis of activations for each experimental condition, relative to rest periods, in the area V5/MT. One-sample t-tests were again used to compare the activations for each condition with activations during rest periods, separately in the blind and the sighted group. Bonferroni correction was used to correct the results across all conditions, for each participant group separately. The differences across experimental conditions and groups were tested with mixed ANOVA. Bonferroni correction was used in pairwise comparisons, when applicable.

## Acknowledgements

This work was supported by a National Science Center Poland grant (2020/37/B/HS6/01269) and a Polish National Center for Academic Exchange fellowship (BPN/SEL/2021/1/00004) to Ł.B.

## Author contributions

M.U., A.C. and Ł.B. conceptualized and designed the study; M.U. and M.P. collected the data; M.U. and Ł.B. performed the data analyses; M.U. and Ł.B. wrote the manuscript; A.C. revised the manuscript.

