## Supplementary Materials for "Neural representation of nouns and verbs in congenitally blind and sighted individuals"

**Supplementary Figures**

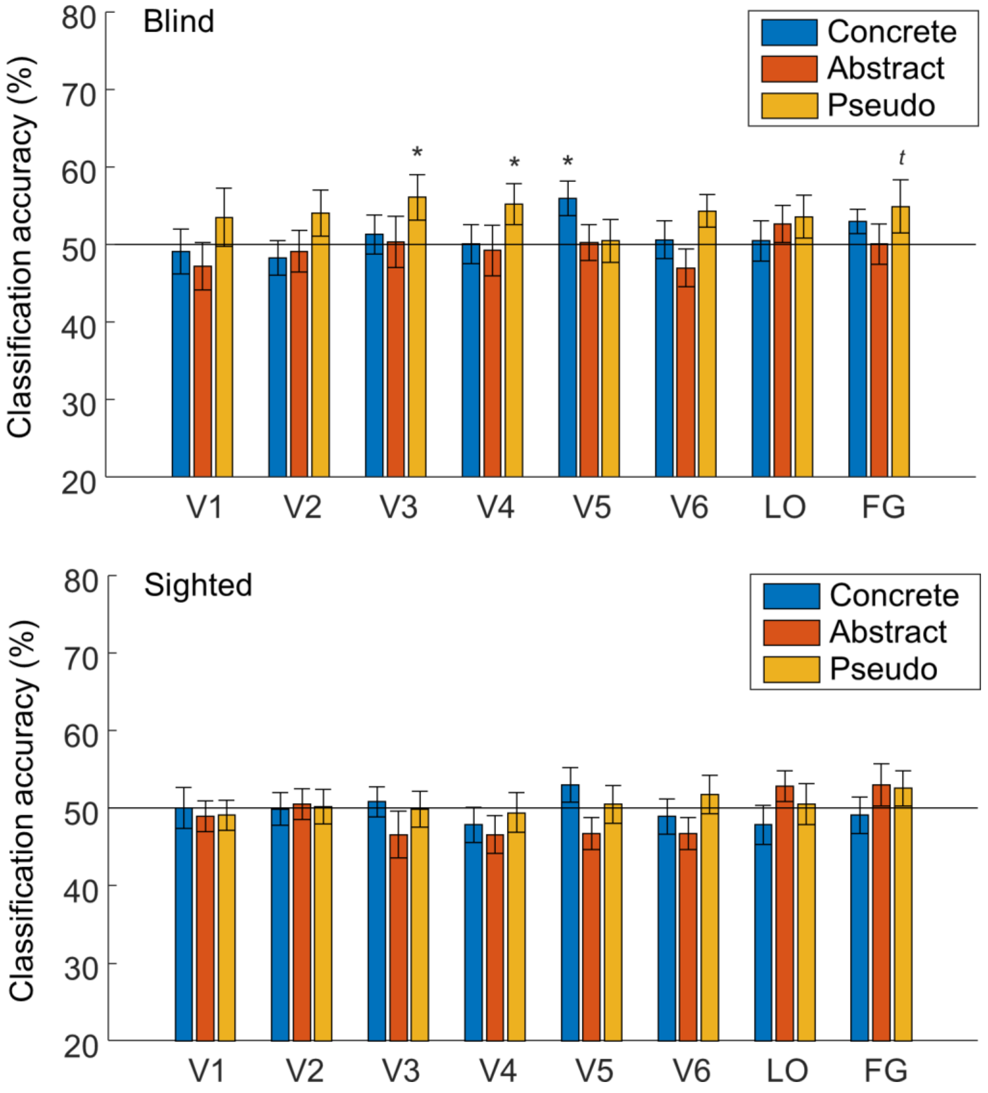


**Figure S1. Classification of activity patterns for nouns and verbs from specific semantic categories in all visual areas included in the study.** Results of support vector machine classification of activity patterns for noun blocks and verb blocks, performed separately for concrete, abstract, and pseudo word categories, in the visual areas in congenitally blind and sighted participants. * $p < 0.05$, $^{t}$ $p = 0.051$, corrected for multiple comparisons using Bonferroni correction. Error bars represent the standard error of the mean. The black lines indicate the chance classification level.

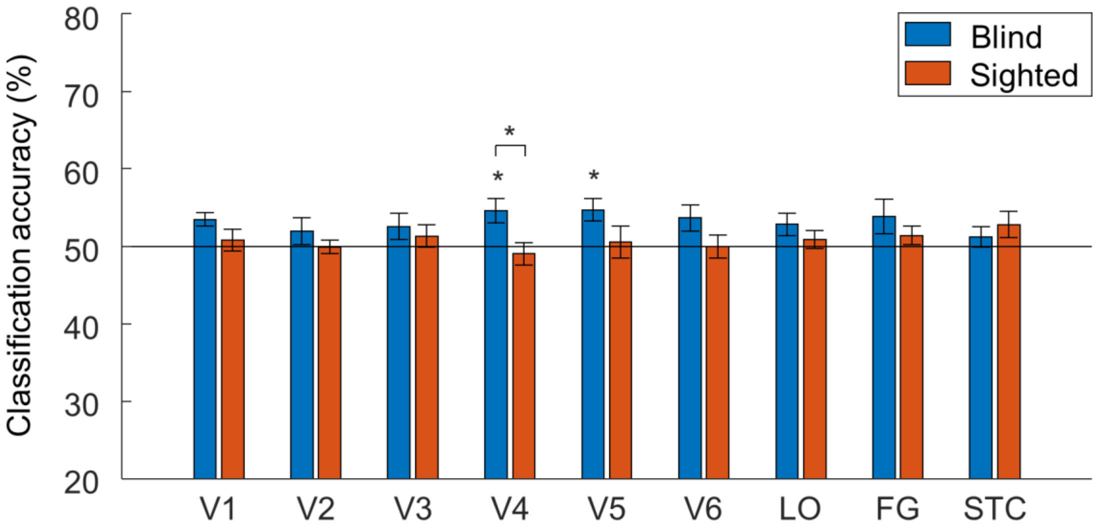


**Figure S2. The above-chance classification of activity patterns for concrete and abstract words in areas V4 and V5/MT in congenitally blind individuals** Results of support vector machine classification of activity patterns for concrete word blocks and abstract word blocks in the visual areas and the superior temporal cortex (STC) in congenitally blind and sighted participants. * $p < 0.05$, corrected for multiple comparisons using Bonferroni correction. Error bars represent the standard error of the mean. The black line indicates the chance classification level.

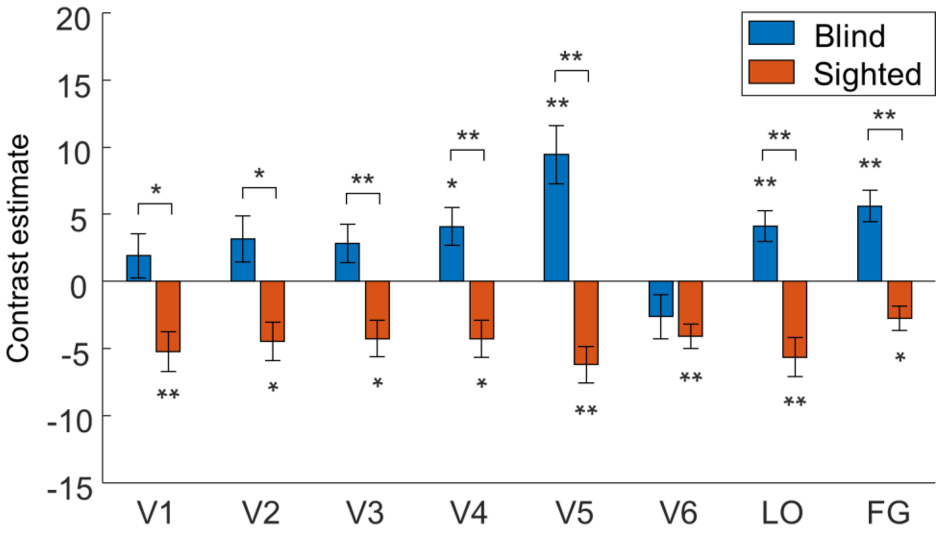


**Figure S3. Spoken words activated the visual cortex in congenitally blind individuals but deactivated this region in sighted individuals.** Average responses to all spoken words and pseudowords, compared to activations during rest periods, in the congenitally blind and the sighted participants. * $p < 0.05$, ** $p < 0.01$, corrected for multiple comparisons using Bonferroni correction. Error bars represent the standard error of the mean.

**Supplementary Tables**

**Table S1.** The results of the behavioral experiment in which sighted participants (n = 16) performed morphological transformations of words and pseudo words used in the fMRI experiments. The task was the same as in the fMRI experiment, but the participants were asked to produce overt responses, which were recorded and analyzed. Response times (from the onset of word presentation to the onset of response) are in milliseconds. Standard deviations (SDs) are reported in parentheses.

| Word transformations – mean reaction times (SD) | Abstract | Concrete | Pseudo |
|---|---|---|---|
| Nouns | 1276 (86) | 1250 (68) | 1402 (77) |
| Verbs | 1253 (116) | 1233 (125) | 1402 (174) |

**Table S2.** The linguistic properties of words and pseudo words used in the fMRI experiment. The results are presented at the word category level – averages (and SDs) are presented. The word frequencies are quantified in a form of Zipf scores.

| Word linguistic properties | Abstract nouns | Abstract verbs | Concrete nouns | Concrete verbs | Pseudo nouns | Pseudo verbs |
|---|---|---|---|---|---|---|
| Frequency – presented word (Zipf) | 4,115 (0,307) | 3,925 (0,704) | 4,091 (0,459) | 4,033 (0,572) | - | - |
| Frequency – target word (Zipf) | 3,506 (0,604) | 3,46 (0,616) | 3,622 (0,664) | 3,512 (0,59) | - | - |
| Syllables – presented word | 2,25 (0,608) | 2,29 (0,464) | 2,25 (0,676) | 2,17 (0,381) | 2,29 (0,464) | 2,25 (0,442) |
| Syllables – target word | 2,75 (0,794) | 2,79 (0,588) | 2,83 (0,702) | 2,71 (0,69) | 2,79 (0,658) | 2,75 (0,608) |

**Table S3.** The ratings for words used in the fMRI experiment provided by a group of congenitally blind participants (n = 15). The participants were asked to rate words on three 1 to 7 scales. The results are presented at the word category level – averages (and SDs) are presented.

| Word ratings – blind participants | Concreteness | Imageability | Movement connotations |
|---|---|---|---|
| Abstract nouns | 3,17 (0,61) | 3,43 (0,77) | 2,33 (0,68) |
| Abstract verbs | 3,27 (0,72) | 3,74 (0,58) | 3,61 (1,01) |
| Concrete nouns | 6,82 (0,23) | 6,72 (0,33) | 1,58 (0,84) |
| Concrete verbs | 6,26 (0,47) | 6,43 (0,32) | 6,22 (0,69) |

**Table S4.** The ratings for words used in the fMRI experiment provided by a group of sighted participants (n = 46). The participants were asked to rate words on three 1 to 7 scales. The results are presented at the word category level – averages (and SDs) are presented.

| Word ratings - sighted participants | Concreteness | Imageability | Movement connotations |
|---|---|---|---|
| Abstract nouns | 3,26 (0,57) | 3,34 (0,5) | 2,43 (0,5) |
| Abstract verbs | 3,49 (0,44) | 3,57 (0,5) | 3,44 (0,75) |
| Concrete nouns | 6,72 (0,21) | 6,78 (0,17) | 1,7 (0,6) |
| Concrete verbs | 5,44 (0,29) | 5,77 (0,35) | 5,23 (1,13) |

**Appendix 1.** The approximate English translations of all words used in the fMRI experiment.

| Word category | Word (English translation) |
|---|---|
| Abstract nouns | issue |
| Abstract nouns | motive |
| Abstract nouns | outcome |
| Abstract nouns | secret |
| Abstract nouns | deadline |
| Abstract nouns | habit |
| Abstract nouns | decision |
| Abstract nouns | harm |
| Abstract nouns | opinion |
| Abstract nouns | confidence |
| Abstract nouns | weakness |
| Abstract nouns | property |
| Abstract nouns | order |
| Abstract nouns | trick |
| Abstract nouns | project |
| Abstract nouns | diagram |
| Abstract nouns | style |
| Abstract nouns | talent |
| Abstract nouns | possibility |

| | |
|---|---|
| Abstract nouns | promise |
| Abstract nouns | try |
| Abstract nouns | difference |
| Abstract nouns | loose |
| Abstract nouns | ability |
| Abstract verbs | to defend |
| Abstract verbs | to remember |
| Abstract verbs | to assume |
| Abstract verbs | to spend |
| Abstract verbs | to find a way |
| Abstract verbs | to call for |
| Abstract verbs | to exist |
| Abstract verbs | to belong |
| Abstract verbs | to try |
| Abstract verbs | to cause |
| Abstract verbs | to make something happen |
| Abstract verbs | to doubt |
| Abstract verbs | to name |
| Abstract verbs | to fulfill |
| Abstract verbs | to lose |

| | |
|---|---|
| Abstract verbs | to be able to |
| Abstract verbs | to blame |
| Abstract verbs | to force someone to something |
| Abstract verbs | to finish |
| Abstract verbs | to serve |
| Abstract verbs | to lose |
| Abstract verbs | to pretend |
| Abstract verbs | to believe |
| Abstract verbs | to begin |
| Concrete nouns | carpet |
| Concrete nouns | button |
| Concrete nouns | flower |
| Concrete nouns | container |
| Concrete nouns | towel |
| Concrete nouns | table |
| Concrete nouns | hand |
| Concrete nouns | newspaper |
| Concrete nouns | cable |
| Concrete nouns | rock |
| Concrete nouns | women |

| | |
|---|---|
| Concrete nouns | dress |
| Concrete nouns | pen |
| Concrete nouns | diploma |
| Concrete nouns | computer |
| Concrete nouns | chicken |
| Concrete nouns | writing |
| Concrete nouns | telephone |
| Concrete nouns | battery |
| Concrete nouns | bottle |
| Concrete nouns | book |
| Concrete nouns | kitchen |
| Concrete nouns | plate |
| Concrete nouns | face |
| Concrete verbs | to run |
| Concrete verbs | to touch |
| Concrete verbs | to talk |
| Concrete verbs | to scream |
| Concrete verbs | to throw |
| Concrete verbs | to hear |
| Concrete verbs | to exercise |

| | |
|---|---|
| Concrete verbs | to catch |
| Concrete verbs | to talk |
| Concrete verbs | to write |
| Concrete verbs | to go across |
| Concrete verbs | to get up |
| Concrete verbs | to walk |
| Concrete verbs | to sit |
| Concrete verbs | to sing |
| Concrete verbs | to run away |
| Concrete verbs | to sit |
| Concrete verbs | to go out |
| Concrete verbs | to drive |
| Concrete verbs | to kick |
| Concrete verbs | to open |
| Concrete verbs | to go |
| Concrete verbs | to jump |
| Concrete verbs | to dance |
